# Anoctamin 9/TMEM16J is a Cation Channel Activated by cAMP/PKA Signal

**DOI:** 10.1101/188243

**Authors:** Hyungsup Kim, Hyesu Kim, Jesun Lee, Byeongjun Lee, Heeryang Kim, Jooyoung Jung, Mi-Ock Lee, Uhtaek Oh

## Abstract

Anoctamins are membrane proteins that consist of 10 homologs. ANO1 and ANO2 are anion channels activated by intracellular calcium that meditate numerous physiological functions. ANO6 is a scramblase that redistributes phospholipids across the cell membrane. However, the others are not well characterized. We found ANO9/TMEM16J is a cation channel activated by a cAMP-dependent PKA. Intracellular cAMP activated robust currents in whole-cells expressing ANO9 and inhibited by PKA blockers. A cholera toxin and purified PKA also activated ANO9. The cAMP-induced ANO9 currents were permeable to cations. The mutation of a possible phosphorylation site at Ser245 elicited a block of the cAMP-dependent activation. High levels of *Ano9* transcripts were found in intestines. Human intestinal SW480 cells showed cAMP-dependent currents. We conclude that ANO9 is a cation channel activated by the cAMP/PKA pathway and could play a role in intestine function.

## INTRODUCTION

The Anoctamin/TMEM16 family consists of transmembrane proteins in 10 isoforms, ranging from ANO1/TMEM16A to ANO10/TMEM16K. Anoctamins are expressed in numerous major tissues and are thought to mediate various physiological functions. The best known anoctamin gene is *Ano1*, which is a Cl^-^ channel activated by Ca^2+^ (Caputo et al, 2008; Schroeder et al, 2008; Yang et al, 2008). ANO1 is known to mediate transepithelial fluid movements such as salivation in the salivary glands, mucin secretion in the airway and Cl^-^ and fluid secretion in the intestine (Huang et al, 2012a; Jang & Oh, 2014; Namkung et al, 2011; Ousingsawat et al, 2009; Romanenko et al, 2010). ANO1 is also highly expressed in small-diameter dorsal-root ganglion neurons, implicating its role in nociception as a heat sensor (Cho et al, 2012). In addition, ANO1 plays an important role in controlling smooth muscle contraction (Bulley et al, 2012) and pacemaking activity in the intestine (Cobine et al, 2017; Hwang et al, 2009). More importantly, ANO1 has been implicated in tumorigenesis (Britschgi et al, 2013; Duvvuri et al, 2012; Jia et al, 2015) and benign prostate hyperplasia (Cha et al, 2015).

ANO2 is a Cl^-^ channel activated by Ca^2+^ and is thus also considered a Ca^2+^-activated Cl^-^ channel (Huang et al, 2012a; Pifferi et al, 2009). ANO2 has physiological functions distinct from ANO1; it controls sensory transduction and synaptic plasticity in the central nervous system (Huang et al, 2012b; Zhang et al, 2015) as well as olfactory transduction and phototransduction (Billig et al, 2011; Dauner et al, 2013; Keckeis et al, 2017; Pietra & Dibattista, 2016; Stephan et al, 2009). In addition, ANO2 is also involved in smooth muscle contraction (Bernstein et al, 2014; Forrest et al, 2010).

Unlike ANO1 and ANO2, ANO6/TMEM16F has dual functions. ANO6 is a small conductance calcium activated cation channel (SCAM) that is permeable to divalent ions (Yang et al, 2012). Strikingly, ANO6 is known to be a scramblase that disrupts polarized phospholipids in the plasma membrane. The polarized phospholipids attract immunological signals necessary for activation of T lymphocytes (Hu et al, 2016; Xu et al, 2008; Zhang et al, 2011). Mutations in *Ano6* cause a rare bleeding disease, Scott syndrome (Castoldi et al, 2011; Suzuki et al, 2010). Thus, ANO6 is both a channel and an enzyme. Some scramblase activity has also been observed in ANO4, ANO8 and ANO9.

ANO5 is involved in skeletomuscular function; its mutations cause gnathodiaphyseal dysplasia, an autosomal dominant inherited bone disorder (Tran et al, 2014). However, its biophysical properties and mechanism of activation are unknown.

Although the functional roles of some genes in the Anoctamin family are well studied, the role of ANO9/TMEM16J remains poorly understood (Picollo et al, 2015). ANO9 is found in the human nasal and colonic epithelium and expressed in the respiratory, digestive, skeletal and integumentary systems during development (Rock & Harfe, 2008). ANO9 is also implicated in the metastasis of colorectal cancer (Li et al, 2015). Despite its expression pattern and a possible role in colorectal tumorigenesis, the function and activation mechanism of ANO9 is not known. The present study aimed to determine if ANO9 is a channel and, if so, how it is activated. Surprisingly, we found that ANO9 is a cation channel activated by the cAMP/PKA pathway.

## RESULTS

### cAMP activates ANO9

To determine if ANO9 is a channel, we expressed mouse ANO9 tagged on the C terminus with enhanced green fluorescence protein (eGFP) for visual identification in HEK 293T cells. The eGFP was largely localized in the plasma membrane (Fig. 1A). Upon whole-cell formation in the HEK 293T cells transfected with *Ano9-eGFP* (ANO9/HEK cells), robust inward currents (65.9 ± 6.99 pA/pF, n = 36) were observed with E_hold_ = −60 mV when the pipette solution contained 100 μM cAMP. The bath and pipette solutions contained 140 mM NaCl and 140 mM KCl, respectively. These currents were not observed in whole cells without cAMP in the pipette. In addition, the cAMP-evoked currents were completely blocked by the treatment of the cells with PKA blockers (20 μM H-7 or 20 μM H-89; Fig. 1B & 1C). cGMP in the pipettes (100 nM) failed to evoke currents in ANO9/HEK cells (Fig. 1D). The cAMP-evoked currents in the ANO9/HEK cells began to develop 5 s to a few minutes after the formation of whole-cells. This time lag suggests a possible indirect signaling pathway for the activation by cAMP

**Figure 1.**
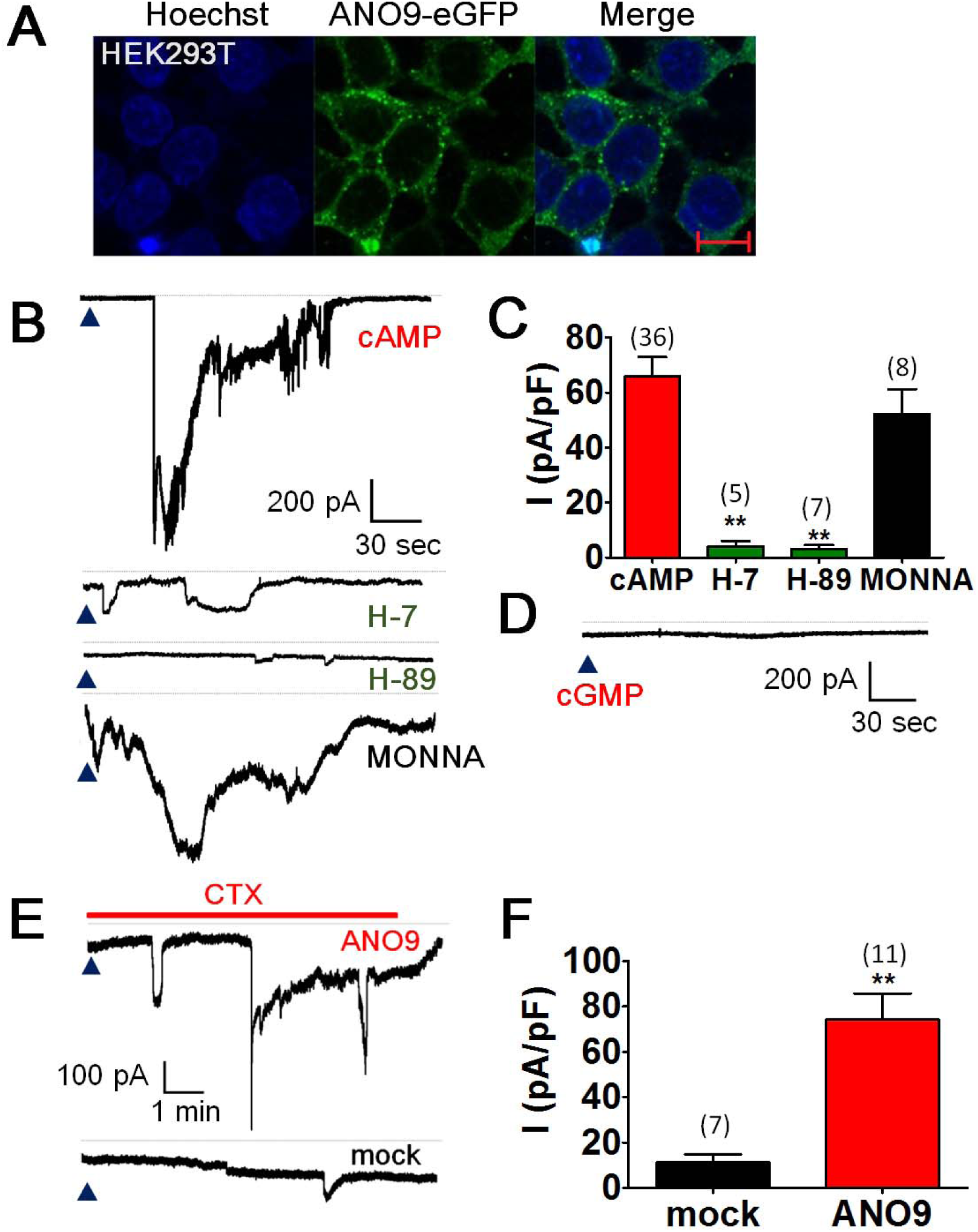
Intracellular cAMP activates ANO9. (A) The overexpression of mANO9-pEGFP-N1 in the HEK293T cells with Hoechst 33342 staining (left). (B) Representative traces of whole-cell currents of HEK293T cells transfected with Ano9. The bath solution contained (in mM) 140 NaCl, 2 MgCl_2_ and 2 CaCl_2_ buffered with 10 HEPES, and the pipette solution contained 140 KCl, 2 MgCl_2_, 2 ATP, 0.3 GTP buffered with 10 mM HEPES. cAMP (100 μM) was added to the pipette solution. E_hold_ = −60 mV. Note that intracellular cAMP evoked robust inward currents in the ANO9-HEK cells but not in the H-7 and H-89 pre-treated cells. The pretreatment of an ANO1-antagonist, MONNA, failed to block the cAMP-induced ANO9 current. The dashed lines represent zero current. Arrow head represents the time for forming a whole cell. (C) Summary of the cAMP-induced whole-cell currents of control (cAMP) and the H-7, H-89, MONNA pre-treated ANO9-HEK cells. The error bars represent S.E.M. ** p < 0.01, one way ANOVA followed by Tukey’s post-hoc test. Numbers in parenthesis represent the experimental numbers. (D) A representative trace of whole-cell currents of ANO9-HEK cells with 100 μM cGMP in the pipette solution. (E) The cholera toxin (CTX) applied to the bath evoked whole-cell inward currents in ANO9-HEK cells but not in mock-transfected HEK293T cells. (F) Summary of the CTX-induced currents in ANO9-HEK cells.

Cholera toxin is an oligomeric protein complex secreted by the bacterium *Vibrio cholerae* that leads to the direct activation of adenylyl cyclase, resulting in the overproduction of cAMP (Gabriel et al, 1994). If ANO9 is activated by cAMP and its downstream signal, then the application of cholera toxin should activate ANO9. Indeed, the application of cholera toxin to the bath solution evoked robust currents with variable time lags in ANO9/HEK cells. However, the cholera toxin-induced currents were not observed in the mock-transfected HEK 293T cells (Fig. 1E & 1F).

### PKA activates ANO9

We then tested whether cAMP alone or with its downstream signal, PKA, activated ANO9. To do this, membrane patches were isolated from ANO9/HEK cells with an inside-out patch configuration. Recordings were made in a pipette solution containing 140 mM NaCl and 2 mM CaCl2 and a bath solution containing 140 mM KCl at E_hold_ = −60 mV. When solutions containing 1 mM cAMP, 2 mM ATP or cAMP+ATP were applied to the inside-out patches, no appreciable currents were observed (Fig. 2A). However, when purified PKA (10 μg/reaction, 0.7 units/μg) along with ATP and cAMP was applied to isolated membrane patches of ANO9/HEK cells, large macroscopic currents were observed with a prolonged time lag (Fig. 2A). These results now suggest that ANO9 is activated by PKA activity.

**Figure 2.**
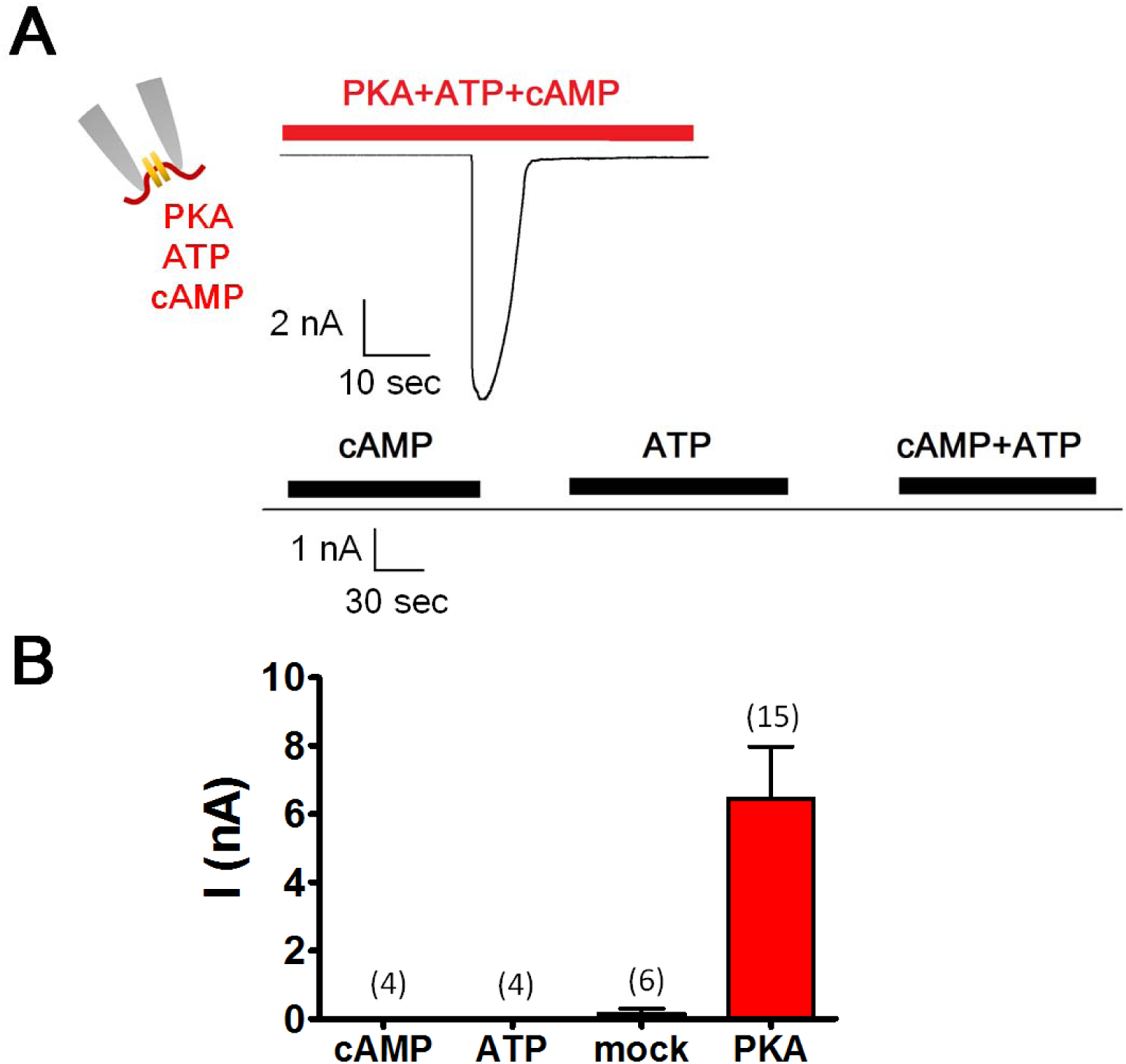
The phosphorylation by PKA activates ANO9. (A) Representative traces of macroscopic currents of inside-out membrane patches from ANO-HEK cells. The application of purified PKA proteins (7 units/ml) together with 100 μM cAMP and 2 mM ATP to the bath evoked a robust current (upper) whereas the application of cAMP and ATP alone failed to evoke the macroscopic currents. (B) Summary of the PKA-evoked currents in ANO9-HEK cells in inside-out patch configuration.

### ANO9 is a cation channel

The cAMP-evoked currents in ANO9/HEK cells were cationic because they were not observed in the bath solution containing 140 mM N-methyl D-glucamine (NMDG)-Cl (Fig. 3A & 3B). Ion selectivity was determined by a shift in reversal potential in whole cells in which extracellular KCl solution was changed to 70 and 210 mM. The pipette solution contained 140 mM KCl. The current-voltage (I–V) relationship was obtained to measure the reversal potential. Voltage ramps from −100 mV to +100 mV in 100 ms durations were applied. When the bath KCl concentration, initially 140 mM, was changed to 210 mM and 70 mM, the reversal potentials were changed to +6.29 ± 0.82 mV and −11.46 ± 1.09 mV (n = 8), respectively (Fig. 3C). The reversal potentials were plotted as a function of extracellular KCl concentration on a semi-logarithmic scale. A line was fitted to +37.2 mV/decade, which is relatively close to +58 mV/decade of the Nernst equation when the major carrier charge is a cation. The relative permeability ratio of K^+^ and Cl^-^ (P_Cl_/P_K_) calculated by Goldman-Hodgkin-Katz equation (Hille, 2001) is 0.237 ± 0.034, suggesting a weak permeability to Cl^-^. Thus, these results clearly suggest that ANO9 exhibits a large preference for cations.

**Figure 3.**
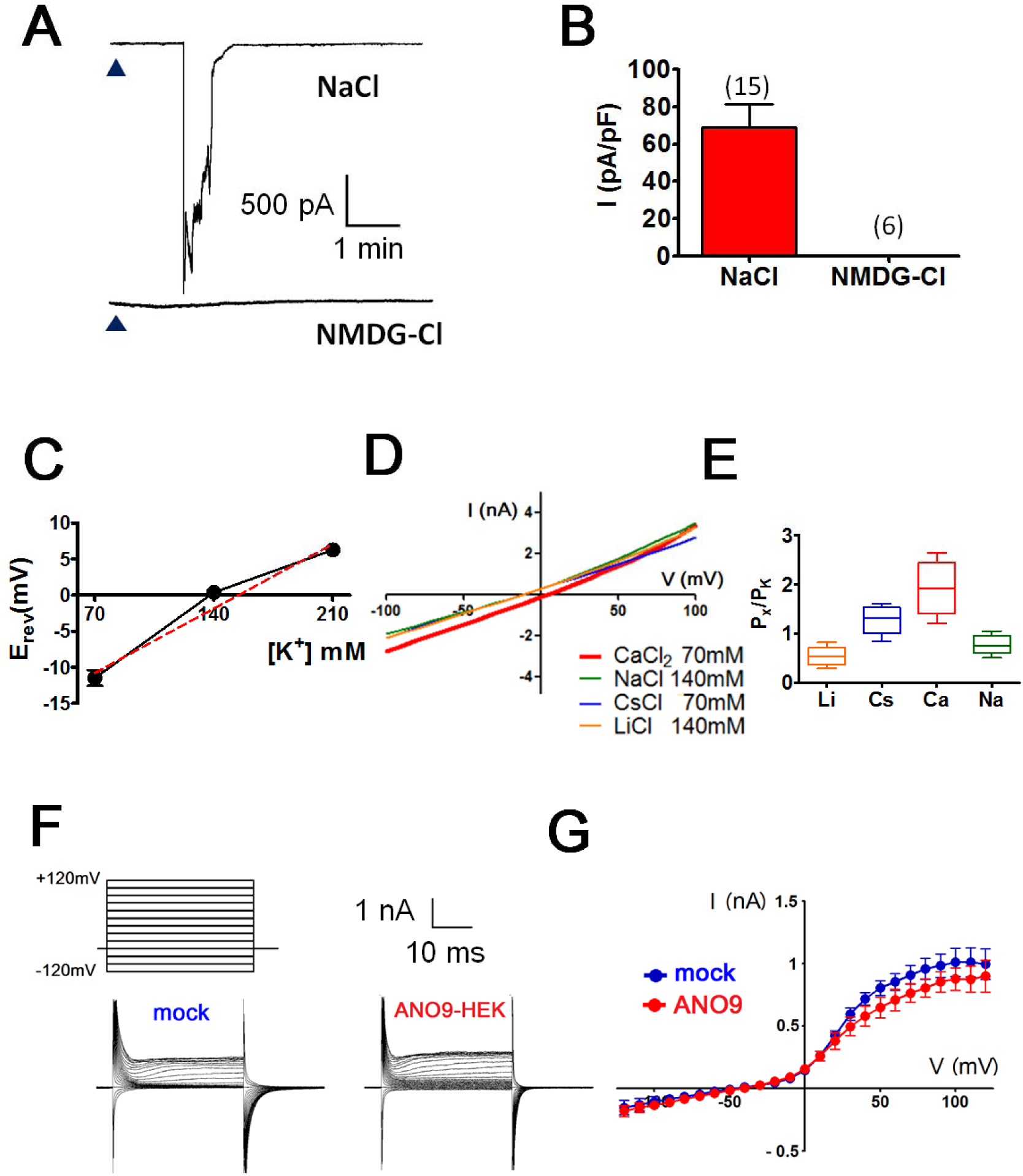
ANO9 is a cation channel. (A) Whole-cell currents of ANO9-HEK cells activated by 100 μM cAMP in 140 mM NaCl (upper) but not in 140 mM NMDG-Cl (lower) bath solutions. (B) The average current densities of ANO9-HEK cells activated by the intracellular cAMP at NaCl or NMDG-Cl solution condition. (C) Reversal potentials in different KCl concentration gradients between pipette and bath solutions. Reversal potentials were measured under bi-ionic conditions where the pipette contained 140 mM KCl whereas the bath contained 70, 140 or 210 mM KCl. n = 8 (D-E) Cation selectivity of cAMP-induced ANO9 currents. Representative I-V curves (D) and the permeability ratios (P_X_/P_K_) (E) of ANO9 currents. Whole cells were formed with the pipette contained 140 mM KCl. Current-voltage relationship were obtained after the bath solution was changed to 70 mM CaCl_2_, 140 mM NaCl, 70 mM CsCl, and 140 mM LiCl. The permeability ratios, P_X_/P_K_ was calculated using the Goldman-Hodgkin-Katz equation. (F-G) Whole-cell currents (F) and the I-V curves of ANO9-HEK cells elicited by voltage steps. Voltage steps from −120 to +120 mV in 20 mV increment were applied to whole cells of HEK cells transfected with vector alone (mock) or Ano9 (ANO9-HEK).

We then explored the selectivity among cations by estimating the permeability ratios after replacing the 140 mM KCl solution in the bath with a solution of 70 mM CaCl2 and CsCl and 140 mM of NaCl and LiCl. The pipette solution contained 140 mM KCl. The reversal potentials were −2.07 ± 1.56, −7.83 ± 1.25, −5.42 ± 1.43 and −12.6 ± 2.96 mV when the bath K^+^ solution was changed to Ca^2+^, Cs^+^, Na^+^, and Li^+^, respectively (Fig. 3D). The relative permeability ratios (P_X_/P_K_) ranged from 0.54 to 1.87 (Fig. 3E), suggesting that ANO9 discriminated poorly among cations but was more permeable to Ca^2+^ than to monovalent cations.

Because both ANO1 and ANO2 are known to be voltage-activated (Cenedese et al, 2012; Xiao et al, 2011), we investigated if ANO9 was also activated by voltage alone. We increased the voltage in 20 mV steps from −120 mV to 120 mV and applied it to the whole cells of mock-and ANO9-transfected HEK cells. The I–V curves of ANO9-transfected cells were outwardly rectifying (Fig. 3G). However, there was no difference between the amplitudes and shapes of the I–V curves of mock-and ANO9-transfected cells, suggesting that voltage alone does not activate ANO9.

### Possible PKA phosphorylation site

Because ANO9 was activated by PKA, we identified its phosphorylation site. Five putative PKA phosphorylation consensus sites (R-R/K-X-S/T, R-X-X-S/T, and R-X-S/T) were predicted by a computer program, GPS 2.0 (Xue et al, 2008). These putative sites were the S85, S120, S245, S321 and T412 residues in ANO9. Using mutagenesis, each Ser or Thr residue of the predicted phosphorylation sites was replaced with alanine. Each mutant of ANO9 tagged with eGFP was expressed in HEK293T cells and tested for activation by cAMP. Each ANO9 mutant except T412A was well expressed in the plasma membrane of HEK293T cells (Supplementary Fig. 1). Robust cation currents were observed in HEK293T cells transfected with S85A, S120A and S321A mutants by 100 μM cAMP in the pipette. However, cAMP failed to evoke cation currents in HEK293T cells transfected with S245A mutant of ANO9 (Fig. 4A & 4B).

**Figure 4.**
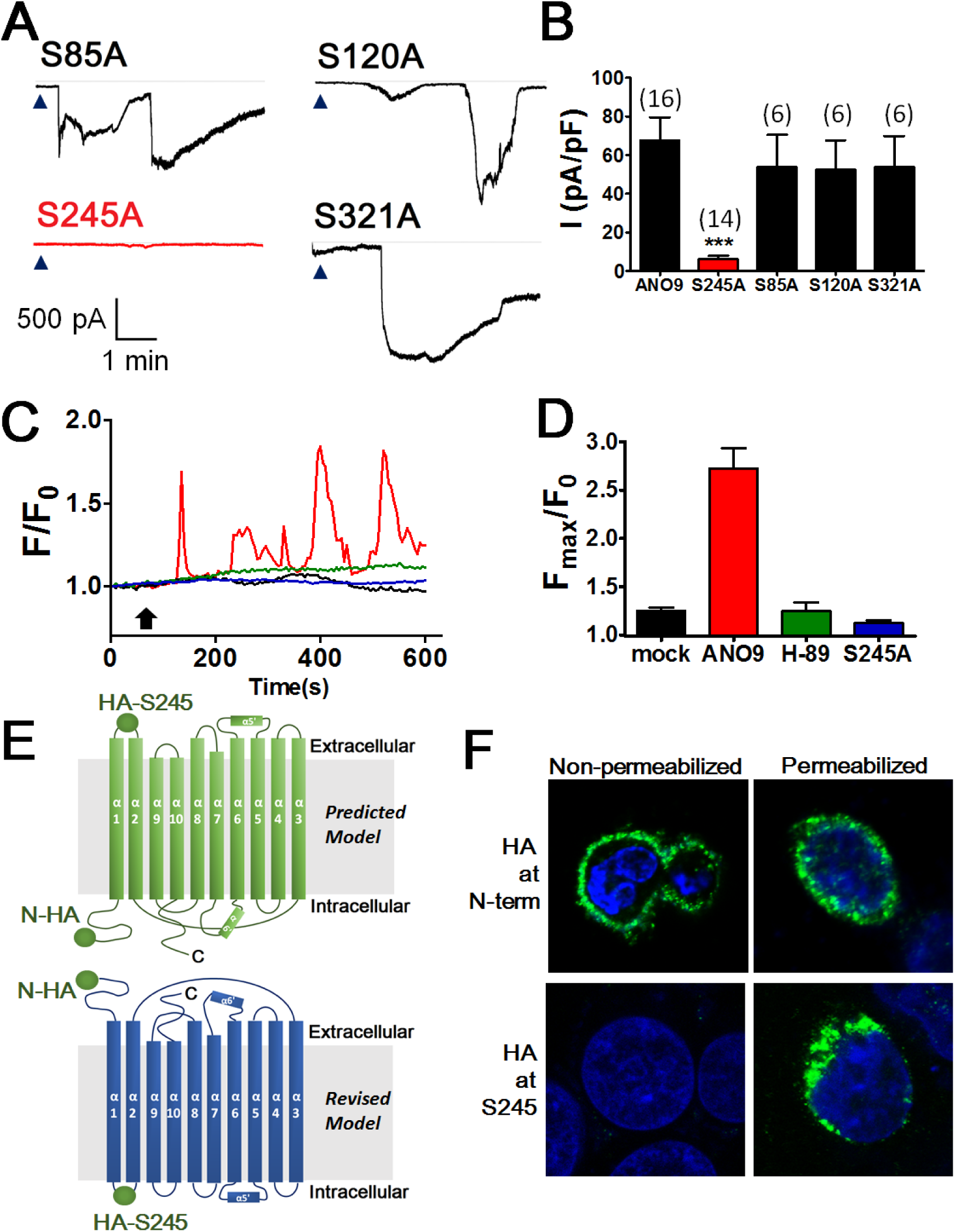
Phosphorylation at Ser245 is essential for ANO9 activity. (A-B) Representative (A) and summary (B) of whole-cell currents of HEK293T cells transfected with ANO9 and its mutants. Putative PKA consensus sites were mutated. Each mutant was transfected to HEK293T cells. Intracellular cAMP evoked robust inward currents in the S85A, S120A and S321A-HEK cells but not in the S245A-HEK cells. *** p < 0.001, one way of ANOVA, Tukey’s *post-hoc* test. (C-D) Representative traces (C) and summary (D) of Ca^2+^ spikes of ANO9-or S245A mutant-transfected HEK293T cells after the application (arrow) of 2 mM dbcAMP Cells were incubated with 1 μM Fluo3-AM. Note that the application of dbcAMP evoked Ca^2+^ spikes in ANO9-HEK cells but not in S245A-HEK or H-89-pretreated ANO9-HEK cells. (E) The predicted topology model of ANO9 extracted from the crystal structure of nhTMEM16 (upper). Bottom image is a revised topology of mANO9. The revised model of mANO9 topology is based on the location of the N-terminus and S245. (F) The immunofluorescent images of hemagglutinin (HA)-tagged Ano9-expressing HEK cells. HA epitope tag was inserted at the N-terminal or at Ser245. The live cells were stained with an antibody for the HA epitope. When the HA was tagged at the N-terminus of ANO9, the HA-tagged ANO9-HEK cells were immunostained with the HA antibody whether the cells were permeabilized by Triton X-100 or not (non-permeabilized), indicating its location at the extracellular side. However, when the HA epitope was inserted next to the S245 residue, the HA-tagged ANO9-HEK cells were immunostained only when the cells were permeabilized with Triton X-100, suggesting its location at the intracellular side.

The activation of each ANO9 mutant by intracellular cAMP was also determined by calcium imaging experiments. The wild-type ANO9-expressing cells displayed irregular calcium signals after the treatment with 2 mM N6,2′-O-dibutyryladenosine 3′,5′-cyclic monophosphate (db-cAMP), a cell-permeable form of cAMP The db-cAMP-induced Ca^2+^ transients were blocked by the pretreatment with H-89 (Fig. 4C). The db-cAMP-evoked calcium transients were not detected in the mock-or S245A mutant-transfected HEK293T cells (Fig. 4C & 4D).

Oddly, the Ser245 residue is located in the extracellular loop of the predicted topology of ANO9 based on the modeling extracted from the crystal structure of nhTMEM16 (Brunner et al, 2014) (Fig. 4E). In this model, the N-and C-termini are present in the cytosolic side of the protein (Predicted model, Fig. 4E). If this is true, the Ser245 is hardly phosphorylated by PKA. However, the actual topology of ANO9 may differ from that predicted by the structure of nhTMEM16. If the Ser245 residue is present in the intracellular side, then the N-and C-termini may be present in the extracellular side of the protein as shown in a revised model in Fig. 4E. To determine if the Ser245 residue is located on the intracellular side, we constructed fusion proteins in which the HA epitope tag was attached to the N-terminus or inserted next to the Ser245 of ANO9. These HA-tagged mutants of ANO9 were transfected to HEK cells. The HA antibody was applied to the extracellular side of the transfected cells and was detected by a fluorescent-tagged secondary antibody to anti-rabbit IgG. As shown in Fig. 4F, the HA tag was detected by the HA antibody applied extracellularly to non-permeabilized as well as permeabilized HEK cells that were transfected with HA-tagged ANO9 in the N-terminus, suggesting that the actual N-terminus of ANO9 is present in the extracellular side. In addition, the HA-tag was detected only in permeabilized cells expressing the ANO9 mutant where the HA tag was inserted next to the Ser245 residue whereas it was not detected by the antibody in non-permeabilized cells (Fig. 4F). These results suggest that the Ser245 residue is located inside the cell.

### Calcium augments the cAMP-induced ANO9 currents

Because ANO1 and ANO2 are known to be activated by intracellular Ca^2+^ we also tested for the activation of ANO9 by Ca^2+^. When 1 μM Ca^2+^ was applied to the pipette, no appreciable whole-cell currents were activated. When 10 μM Ca^2+^ was applied, small currents were observed (Fig. 5A & 5B). When 20 μM cAMP was applied together with 10 μM Ca^2+^ much larger currents than those activated by 10 μM Ca^2+^ alone were observed. These current responses to the co-application of Ca^2+^ and cAMP appeared to be dose dependent because larger currents were observed when 100 and 200 μM cAMP were applied along with 10 μM Ca^2+^ (Fig. 5A & 5B). Although the intracellular Ca^2+^ in a physiological concentration rarely activated ANO9, Ca^2+^ can augment the ANO9 response to cAMP.

**Figure 5.**
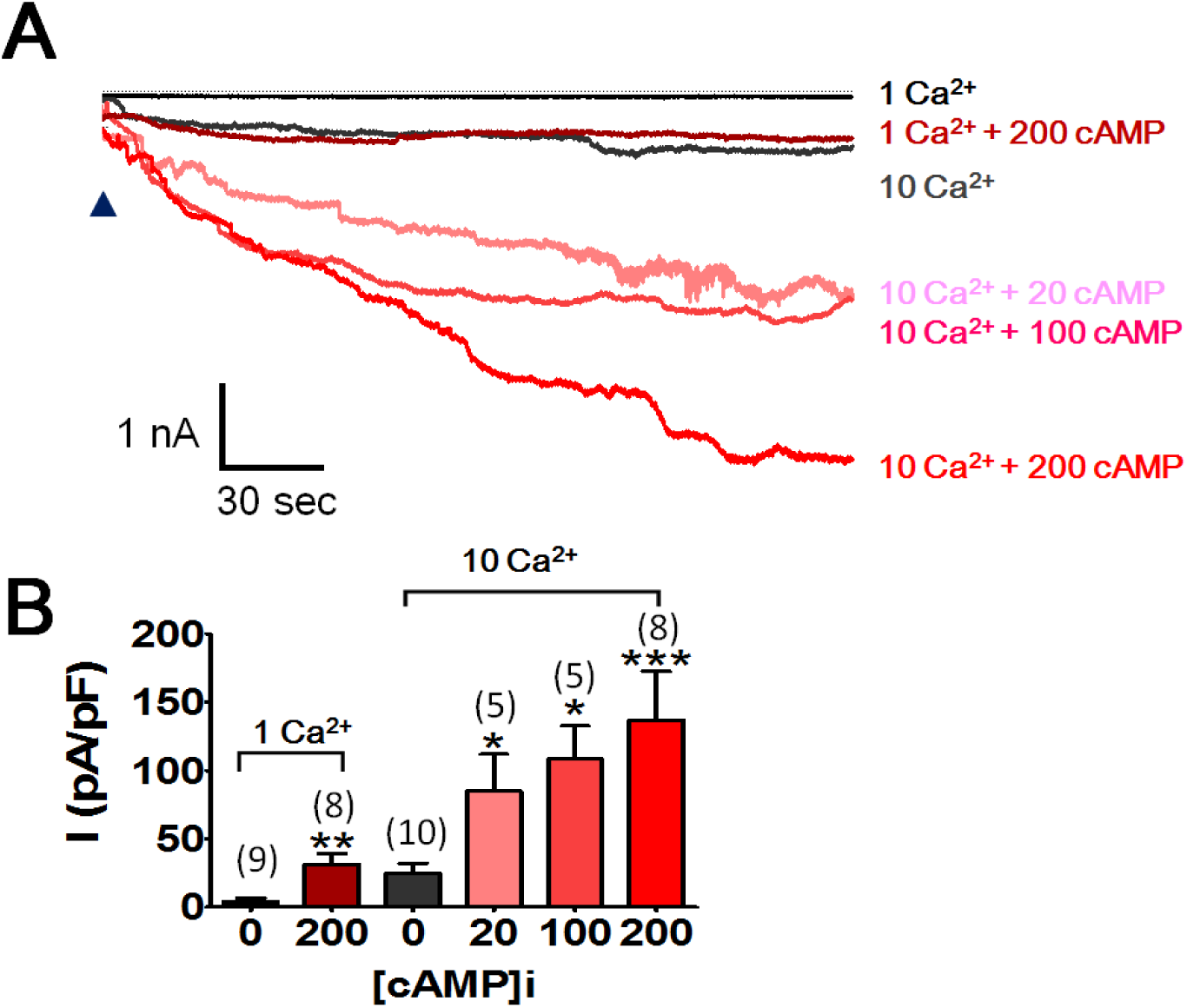
Intracellular calcium augments the cAMP-induced ANO9 activity. (A) Whole-cell currents ANO9-HEK cells activated by various concentrations of intracellular cAMP or/and Ca^2+^. The bath and pipette solutions are same as in Fig. 1. (B) Summary of the cAMP-induced currents at 1 or 10 μM Ca^2+^.* p< 0.01, *** p < 0.001, one way of ANOVA, Tukey’s *post-hoc* test.

### Intracellular sodium inhibits ANO9

Unexpectedly, we also found that a high intracellular concentration of Na^+^ inhibits the activity of ANO9. Intracellular cAMP vigorously activated ANO9 when 0 mM Na^+^ was added to the pipette (intracellular) solution. However, when 50 mM NaCl was added to the pipette solution, the cAMP-evoked whole-cell currents were markedly reduced (Fig. 6A & 6B). In contrast, when 50 mM CsCl was added to the pipette solution, the cAMP-evoked currents were comparable to those of the pipette solution containing 150 mM KCl (Fig. 6A & 6B). These results suggest that high intracellular concentration of Na^+^ inhibited the cAMP-dependent ANO9 activity.

**Figure 6.**
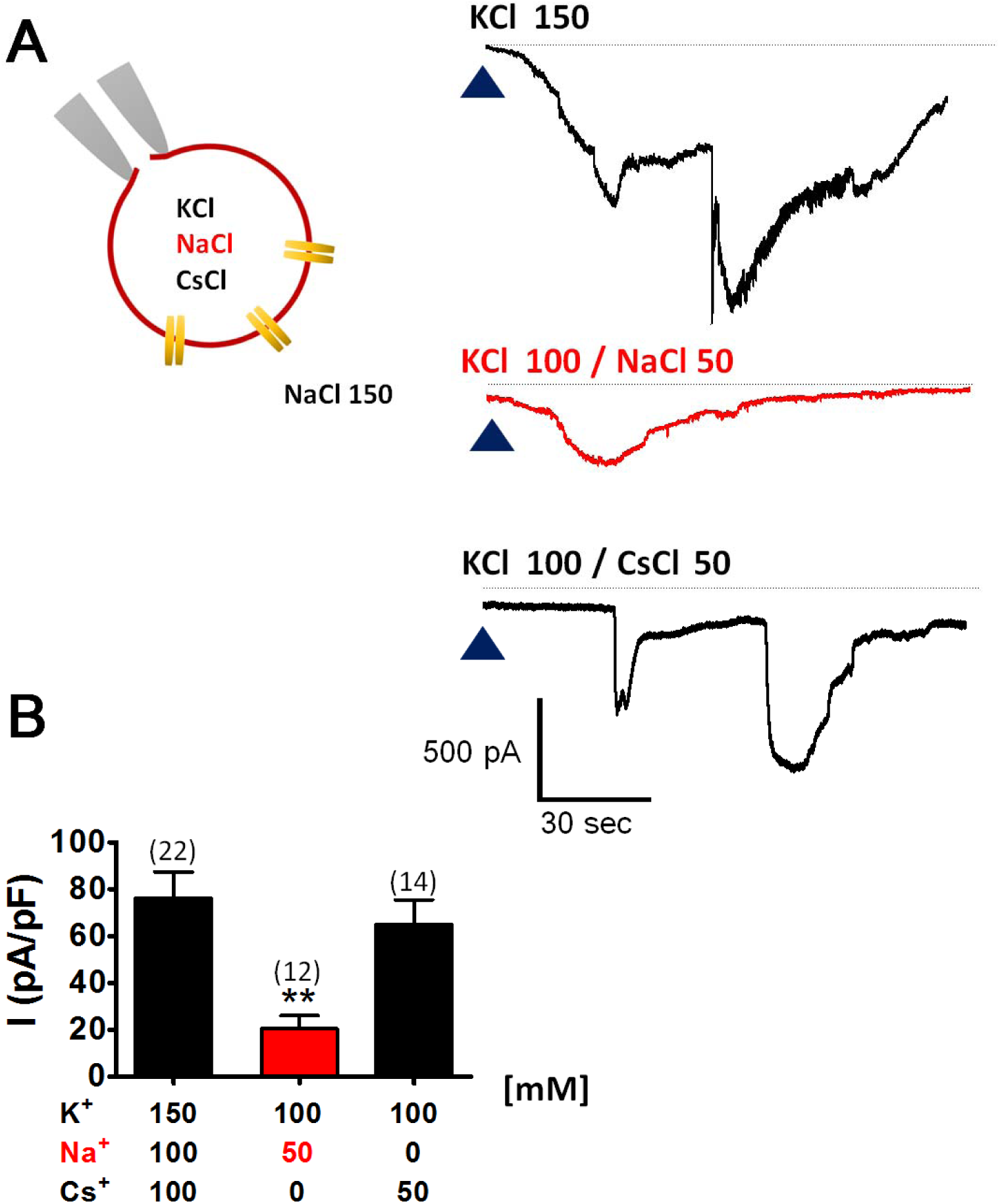
Intracellular sodium inhibits ANO9. (A) cAMP-induced whole-cell currents of ANO9-HEK cells with or without intracellular Na^+^. Concentration of KCl, NaCl or CsCl in the pipette solution was shown in each trace. The bath solution contained 140 mM NaCl. (B) Summary of cAMP-induced currents with different pipette solutions. ** p < 0.01, one way of ANOVA, Tukey’s *post-hoc* test.

### Ano9 is rich in intestines

Tissue distribution of *Ano9* was determined. *Ano9* mRNAs were mainly observed in the digestive system. The small intestine, colon and stomach expressed large number of copies of *Ano9* mRNA. A small amount of *Ano9* mRNA was also detected in the tongue, kidney, eye, lung, and bladder (Fig. 7A).

**Figure 7.**
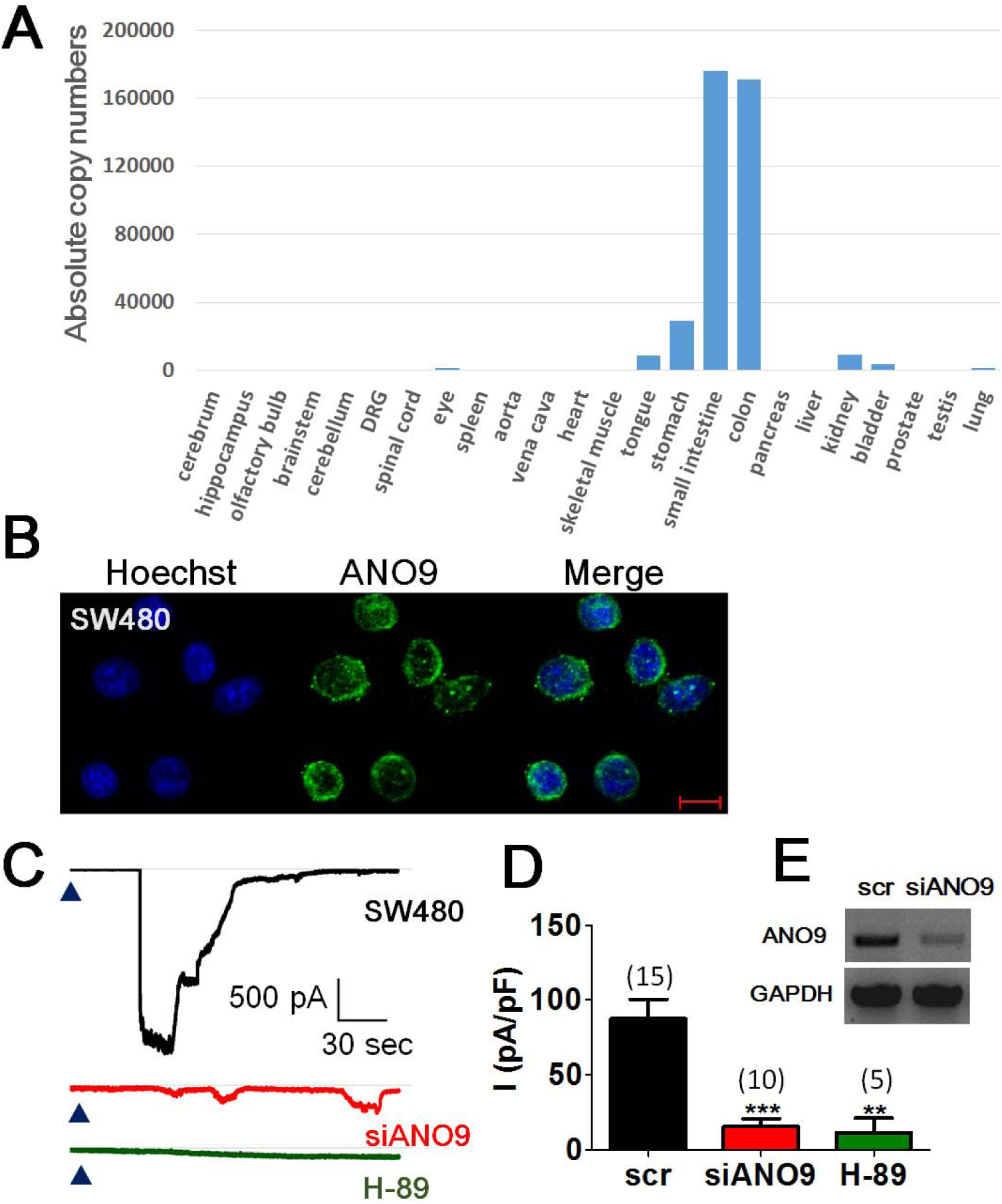
ANO9 is highly expressed in colonic cells. (A) Real-time quantitative PCR analysis of mRNA transcripts encoding ANO9 in various mouse organs. The levels of transcripts were expressed as absolute copy numbers. (B) The immunofluorescence of ANO9 in SW480 cells. Scale bar = 10 μm. (C) Whole-cell currents activated by intracellular cAMP (100 μM) with (upper) or without *Ano9* siRNA treatment (middle). The currents were also blocked by the H-89 pretreatment (lower). (D) Summary of the cAMP-induced currents in the SW480 cells. scr; scrambled siRNA treated, siRNA; *Ano9* siRNA treated; H-89, H-89 treated. (E) Real-time quantitative PCR products of Ano9 in SW480 cells transfected with scrambled or Ano9 siRNAs. PCR products of glyceraldehyde-3-phosphate dehydrogenase (GAPDH) mRNAs were also shown for control.

Consistent with high expression in the colon, ANO9 is known to be expressed in colorectal cancer cells (SW480) (Li et al, 2015). Indeed, the ANO9 specific immunofluorescence was present in the plasma membrane of SW480 cells when probed with ANO9 antibody (Fig. 7B). We therefore determined if cAMP activated the cation currents in these cells. Shown in Fig. 6C, the intracellular application of 100 μM cAMP resulted in large inward currents in 15 out of 20 SW480 cells (Fig. 7C). The cAMP-evoked currents were blocked by co-challenge with 20 μM H-89 in the pipette solution. In addition, a knock-down of *Ano9* in the SW480 cells after transfection with siRNA of *Ano9* markedly reduced the cAMP-evoked currents (Fig. 7C & 7D). These results demonstrate that the native ANO9 in colorectal cells is activated by intracellular cAMP

## DISCUSSION

### Diverse functions of the anoctamin family

The anoctamins have 10 homologs with diverse cellular and physiological functions. ANO1 was first discovered as a calcium-activated chloride channel (Caputo et al, 2008; Yang et al, 2008) and is known to be involved in many physiological functions such as fluid secretion, smooth muscle contraction, nociception, tumorigenesis and cell proliferation (Cho et al, 2012; Duvvuri et al, 2012; Huang et al, 2012a; Jang & Oh, 2014; Katoh & Katoh, 2003; Namkung et al, 2011; Ousingsawat et al, 2009; Romanenko et al, 2010). ANO2 is also known to be a calcium-activated chloride channel with olfaction and learning and memory functions (Billig et al, 2011; Huang et al, 2012b; Neureither et al, 2017; Pietra & Dibattista, 2016; Stephan et al, 2009). ANO3 is known to increase modulation of the activity of the Na^+^-activated K^+^ channel in dorsal-root ganglion neurons, and it also modulates nociception (Huang et al, 2013). ANO5, expressed in muscles and bones, has been implicated in skeletomuscular functions because its mutations cause a skeletomuscular disease, gnathodiaphyseal dysplasia (Tran et al, 2014). Despite their physiological implications, however, the activation mechanisms and other cellular functions of ANO3 and ANO5 are not well characterized. ANO6 was initially characterized as a scramblase that disrupted polarized phospholipids in the plasma membrane (Castoldi et al, 2011; Suzuki et al, 2010). Later, ANO6 was found to be a cation channel activated by Ca^2+^ (Yang et al, 2012). Despite the diverse functions of anoctamins, the activation mechanisms of those other than ANO1, ANO2 and ANO6 are not well understood.

Unlike ANO1 and ANO2, ANO9 was permeated mainly by cations. The ANO9 currents were not blocked by the ANO1 blockers such as MONNA, NPPB, or tannic acid (Fig. 1B & 1C). More importantly, ANO9 was activated by the cAMP/PKA pathway but not by Ca^2+^ or voltage. However, the activation by PKA was augmented by intracellular Ca^2+^. Although ANO9 is known to be expressed in the intestines, its physiological function is largely unknown.

### Role in colorectal cancer cells

ANO9 has been implicated in intestinal functions; notably, its transcription levels were much higher in intestines compared to other tissues (Fig. 7A; (Schreiber et al, 2015; Schreiber et al, 2010)). It is expressed in the colorectal cancer cell line, SW480 cells. We observed cAMP-activated and H-89-reversible currents in the SW480 cells (Fig. 7), and ANO9 has been linked to colorectal cancer (Li et al, 2015). The activity of ANO9 was negatively associated with tumorigenesis in the colon. The expression of ANO9 was higher in non-tumor tissue than in tumorous tissues. ANO9 expression was lower in the recurrent colorectal cancer cells than in non-recurrent colorectal cancer cells. ANO9 overexpression is known to reduce the invasion of cancer cells (Li et al, 2015). Lower levels of ANO9 expression have been associated with poorer prognoses in patients with higher expression levels (Li et al, 2015). Thus, ANO9 appears to play a tumor suppressor role in the intestines, but its precise pathophysiological role needs to be determined.

### Calcium dependence

Ca^2+^ is indispensable for anoctamin functions, and it activates ANO1 and ANO2 (Caputo et al, 2008; Schroeder et al, 2008; Stephan et al, 2009; Yang et al, 2008). Ca^2+^ is also required for the scramblase activity of ANO6 and fungal TMEM16 (Malvezzi et al, 2013; Suzuki et al, 2010; Yang et al, 2012). However, Ca^2+^ was not found to be essential for the activation of ANO9 because physiological concentrations of Ca^2+^ rarely gated ANO9. In addition, cAMP activated ANO9 in Ca^2+^-free conditions, but Ca^2+^ augmented the cAMP-induced ANO9 currents. Thus, the role of Ca^2+^ in activating ANO9 cannot be totally ignored because Ca^2+^ affected the ANO9 activity. Recently, the X-ray crystal structure of nhTMEM16 was determined (Brunner et al, 2014). The Glu residues of ANO1 essential for its activation by Ca^2+^ were located in the subunit cavity of ANO1 in the hydrophobic core of the membrane. The subunit cavity was lined by three glutamates, two aspartates and an asparagine residue in the α-helices 6, 7, and 8 of nhTMEM16 and ANO1. These helices are known to be crucial for the channel activation of ANO1 and its scramblase activity (Brunner et al, 2014). ANO9 has a similar amino-acid sequence in this region (Supplementary Fig. 2): ANO9 has three glutamates, one aspartate and one serine residue in the a-helices 7 and 8. This structural constraint of ANO9 compared to ANO1 predicts the synergistic but not essential role of Ca^2+^ in activating ANO9. Voltage has been found to be another endogenous activator of ANO1 and ANO2, but, analogous to the role of Ca^2+^, ANO9 was not activated by voltage alone (Fig. 3F & 3G). We found that ANO9 added more to the diversity of the functions of the anoctamin family.

### PKA activation

Many of the functions of the cystic fibrosis transmembrane conductance regulator (CFTR), such as the transepithelial Cl^-^ transport in various epithelia, overlap with those of ANO1 mainly because they are both anion channels. Because of this functional overlap, ANO1 has been considered to rescue the defective CFTR functions in cystic fibrosis (Becq et al, 2011; Sondo et al, 2014). CFTR is activated by ATP and PKA and has multiple transmembrane-spanning domains in the cell membrane and a regulatory domain and two nucleotide-binding domains in the cytosolic side (Hwang & Sheppard, 2009). The binding of ATP to the nucleotide-binding domains is known to activate the CFTR while the phosphorylation of the regulatory domain by PKA is a prerequisite for the activation. Unlike the CFTR, ANO9 does not contain nucleotide-binding domains, and therefore ANO9 gating does not require ATP because it is not activated by ATP alone (Fig. 2B). However, whether the phosphorylation of ANO9 at Ser245 by PKA is indispensable or is a prerequisite for its activation remains unclear. ANO9 did not have a bulky regulatory domain with multiple phosphorylation sites as shown in the CFTR, which is the structural basis for the prerequisite requirement for activation. Because only the direct application of PKA along with cAMP and ATP activates ANO9, it is highly likely that ANO9 opens only when it is phosphorylated at Ser245 by PKA. One limitation of this hypothesis is the location of the Ser245 residue. When the location was predicted by the crystal structure of the homologous nhTMEM16 in the fungus *Nectria haematococca*, the Ser245 residue was located in the first extracellular loop. This location is difficult to reconcile as the PKA phosphorylation site of ANO9. However, the crystal structure was obtained with membrane-embedded crystals of nhTMEM16, which have only scramblase activity and not channel activity. Mouse ANO9 also has only 21% sequence homology with nhTMEM16. Therefore, the predicted location of the Ser245 residue in the homology modeling with nhTMEM16 might not represent the actual location in the mouse ANO9. To identify if Ser245 region is located in the intracellular region, we cloned an HA-tagged ANO9. If the N-terminal of ANO9 is in the extracellular region, then the Ser245 will be located in the intracellular loop. In live cell imaging, the HA epitope tagged N-terminal was detected by an HA-antibody without detergent. These results provide strong evidence that the location of the N-terminal of ANO9 is extracellular.

## Material and methods

### Cloning of mouse Ano9 and mutagenesis

Primers were designed using the mouse cDNA sequences of *Ano9* (TMEM16J) from the NCBI database (NM_178381.3). The cDNA encoding *Ano9* has been isolated from the lung of adult C57BL/6J mice. The full-length coding sequence of *Ano9* (Tmem16J; NM_178381.3) was amplified by PCR using site-specific primers: forward primer 5’- GCCACCATGCAGGATGATGAGAGTTCCCAG-3’, reverse primer 5’- GACCGGTCTATACATCCGTGCTCCTGGAAC-3’.

*Ano9* were cloned into pEGFP-N1 to have fusion proteins tagged with EGFP. To express ANO9–GFP fusion protein, a stop codon was deleted from pEGFP-N1-mANO1 using the Muta-Direct site directed mutagenesis kit (iNtRON Biotech). The protein sequence of mouse *Ano9* (NP_848468.2) is

MQDDESSQIFMGPEGDQLPLVEMGSCKPEASDQWDCVLVADLQTLKIQKHAQKQLQFLENLESNGFHFKMLKDQKKVFFGIRADSDVIDKYRTLLMNPEDSGSRDEQSFNIATTRIRIVSFVVNNKLKPGDTFEDLVKDGVFETMFLLHKGEQNLKNIWARWRNMFEPQPIDEIREYFGEKVALYFTWLGWYTYMLVPAAVVGLIVFLSGFALFDSSQISKEICSANDIFMCPLGDHSHRYLRLSEMCTFAKLTHLFDNEGTVLFAIFMALWATVFLEIWKRKRAHEVQSWKLYEWDEEEEEMALELINSPHYKLKDHRHSYLSSTIILILSLFMICLMIGMAHVLVVYRVLAGALFSSLVKQQVTTAVVVTGAVVHYIIIVIMTKVNKYVALKLCKFEESGTFSEQERKFTVKFFILQFFAHFSSLIYIAFILGRINGHPGKSTRLAGLWKLEECHLSGCMMDLFIQMAIIMGLKQTLSNCVEYLCPLLAHKWRLMWASKHGHMSKDPELKEWQRNYYMNPINTFSLFDEFMEMMIQYGFTTIFVAAFPLAPLLALFSNLVEIRLDAIKMVRLQRRLVPRKAKDIGTWLQVLETIGVLAVIANGMVIAFTSEFIPRVVYKYHYGPCRTNR TFTDDCLTNYVNHSLSVFYTKHFNDHSRMEGQENVTVCRYRDYRNEHDYNLSEQFWFILAIRLTFVILFEHFALCIKLIAAWFVPDVPQKVKNEVLQEKYDRIRHRMRFSSRST DV.

All mutants were generated from the wild-type mouse *Ano9* construct tagged with eGFP (pEGFP-N1-mANO9). The amino acid substitution mutants were constructed using a site-directed mutagenesis kit. The construction of mutants was verified with DNA sequencing. To generate HA-tagged m*Ano9* mutants, haemagglutinin (HA) epitopes (YPYDVPDYA) were added to the N-terminus of mAno9 using the HA-pcDNA3.1 vector.

The HA epitopes next to the Ser245 residue in m*Ano9* were introduced by site-directed mutagenesis. The insertion was repeated four times with each set of primers.

1^st^ forward primer: 5’-gatacctgcgactctcatacccatgagatgtgcactttcgc-3’, 1st reverse primer: 5’-gcgaaagtgcacatctcatgggtatgagagtcgcaggtatc-3’,

2^nd^ forward primer: 5’-gcgactctcatacccatacgatgtgagatgtgcactttcg-3’, 2nd reverse primer 5’-cgaaagtgcacatctcacatcgtatgggtatgagagtcgc-3’,

3^rd^ forward primer: 5’-tcatacccatacgatgttccagatgagatgtgcactttcgc-3’, 3rd reverse primer: 5’-gcgaaagtgcacatctcatctggaacatcgtatgggtatga-3’,

4^th^ forward primer: 5’-tcatacccatacgatgttccagattacgctgagatgtgcactt-3’, 4th reverse primer: 5’-aagtgcacatctcagcgtaatctggaacatcgtatgggtatga-3’.

### Cell culture and transfection of Ano9

Cell culture and functional expression of ANO9 or mutants were performed in HEK 293T cells. HEK 293T cells were maintained at 5% CO_2_, and incubated at 37°C in Dulbecco’s modified Eagle’s medium (DMEM) supplemented with 10% fetal bovine serum (FBS), 10 units/mL penicillin and 10 μg/mL streptomycin. To induce ANO9 expression in HEK293T cells, cells were transfected with mAno9 cDNA mixed with the FuGene HD (Roche Diagnostics) transfection reagent. The transfected cells were plated onto glass coverslips. The current responses were recorded 24 to 48 h after transfection.

### Whole-cell and single-channel current recordings

Whole-cell and single-channel current recordings were obtained using either a voltage-clamp technique with an Axopatch 200B amplifier (Molecular Devices) as descried previously (Cho et al, 2012; Lee et al, 2015). Briefly, whole-cell currents were measured after breaking the plasma membrane under the pipette tips. The resistance of the glass pipettes was about 3 mΩ. The junctional potentials were cancelled to zero. Unless otherwise stated, the bath solution contained (in mM) 140 NaCl, 2 CaCl_2_, 2 MgCl_2_ and 10 NaOH-HEPES adjusted to pH 7.2. The pipette solution contained (in mM) 140 KCl, 2 CaCl_2_, 2 MgCl_2_ and 10 KOH-HEPES adjusted to pH 7.2.

For inside-out patch recording, after a giga-seal was formed, the glass pipette was pulled quickly from the cell to isolate a membrane patch. The output of the amplifier was fed to an analog/digital converter (Digidata 1440, Molecular Devices) and stored in a personal computer. The pClamp 10 software was used for I-V curve and other biophysical analysis.

Ca^2+^ was chelated with 10 mM EGTA and 10 mM HEDTA to make the free 1.0 and 10 μM Ca^2+^ in the pipette solutions, respectively. The free Ca^2+^ was calculated using WEBMAXC (http://www.stanford.edu/~cpatton/webmaxcS.htm).

### Immunofluorescent staining

SW480 cells were fixed on glass coverslips in 4% paraformaldehyde for 10 min at room temperature, permeabilized with 0.2% Triton X-100, and incubated with anti-ANO9 antibody (dilution 1:200, LS-C179041, LifeSpan BioSciences, Inc.) overnight at 4°C. The coverslips were washed and incubated with the Alexa Fluor 488-tagged donkey-anti-mouse IgG (1:1000, Molecular Probes, CA). For nuclear staining, SW480 cells and HEK293T cells were incubated with Hoechst 33342 (H3570, Thermofisher Scientific, 1:2,000) after ANO9 immunostaining

For HA staining, HEK293T cells transfected with the HA-tagged Ano9 mutants were incubated with anti-HA antibody (1:100, Cat# H6905, Sigma-Aldrich) for 1 hr at room temperature. The cells were then washed three times and incubated with the Alexa Fluor 488 goat-anti-rabbit IgG (1:200, Life Technologies Cat# A11008) for 30 min at room temperature. Finally, the cells were washed and fixed with 4% PFA. For permeabilized staining, cells were fixed with 4% PFA and treated with 0.2% Triton 100 for 10 min. Then, the cells were incubated with the anti-HA antibody (1:100) for 1 hr, washed three times, and then incubated with the Alexa Fluor 488 goat-anti-rabbit IgG for 30 min at room temperature. After washing, the coverslips were mounted and imaged using a confocal microscopy.

### Small interfering RNA (siRNA) treatment

Two nucleotide oligomer siRNAs targeting mouse ANO9 or scrambled siRNA labelled with Cy3 were provided by Bioneer (Seoul, Korea) transfected into cells using Lipofectamine 2000TM (Invitrogen) 1 day after plating. We transfected the cells with a mixture of the two siRNAs simultaneously for maximal effect. The two siRNAs (Nos. 1152997 and 1152999) are as follows.

1152997 Sense: GUGAACUUCGUUGUCAUGA(dTdT), 1152997 Antisense: UCAUGACAACGAAGUUCAC(dTdT),

1152999 Sense: UCAGCAACUGCGUCGAGUA(dTdT), 1152999 Antisense: UACUCGACGCAGUUGCUGA(dTdT).

### Calcium imaging

ANO9-HEK cells or HEK293T cells transfected with S245A *Ano9* mutant were loaded with Fluo3-AM (Invitrogen) containing 0.1% Pluronic F-127 (Invitrogen). After loading with Fluo3-AM for 40 min, db-cAMP alone or with H89 was applied to the cells. The fluorescence intensities of cells were measured at 488 nm in every 5 s with a confocal microscope (LSM700, Zeiss).

### Real-time quantitative PCR

Major mouse organs were isolated from three 7-week-old mice. The total RNAs from each organ were purified with ethanol precipitation or the Easy Spin™ Total RNA Extraction Kit (iNtROn Biotech) according to the manufacturer’s protocol. The first strand cDNAs were reverse transcribed from the total RNA using the Transcriptor First Strand cDNA Synthesis Kit (Roche). To measure the gene expression level, we performed real-time quantitative PCR (qPCR) in the LightCycler 2.0 system (Roche) using an ANO9 specific universal probe (forward primer: cggactctcctcatgaatcc; reverse primer: tgttcacgacaaagctcaca). The absolute copy numbers per 250 ng of total RNA were calculated using the absolute quantification with the external standard method in the Light Cycler software 4.0.

### Reagents

The Anti-ANO9/TMEM16J antibody (aa81-224) LS-C179041 was purchased from Lifespan Biosciences Inc, We purchased adenosine 3′,5′-cyclic monophosphate (cAMP; A9501), cholera toxin from *Vibrio cholera* (CTX; C8052), N6,2′-O-dibutyryladenosine 3′,5′-cyclic monophosphate (db-cAMP) sodium salt (D0627), protein kinase A from bovine heart (PKA; P5511), H-89 dihydrochloride hydrate (B1427), and H-7 dihydrochloride (I7016) from Sigma-Aldrich. All basic chemicals for the electrophysiology experiments were also purchased from Sigma-Aldrich.

#### Statistical analysis

All results are expressed as means ± standard errors (S.E.). The statistical significances of the differences were determined by a one-way analysis of variance (ANOVA) followed by the Tukey’s post-hoc test for multiple comparisons. Statistical significance was accepted at p values of less than 0.05.

---

**Supplementary Figure 1.**
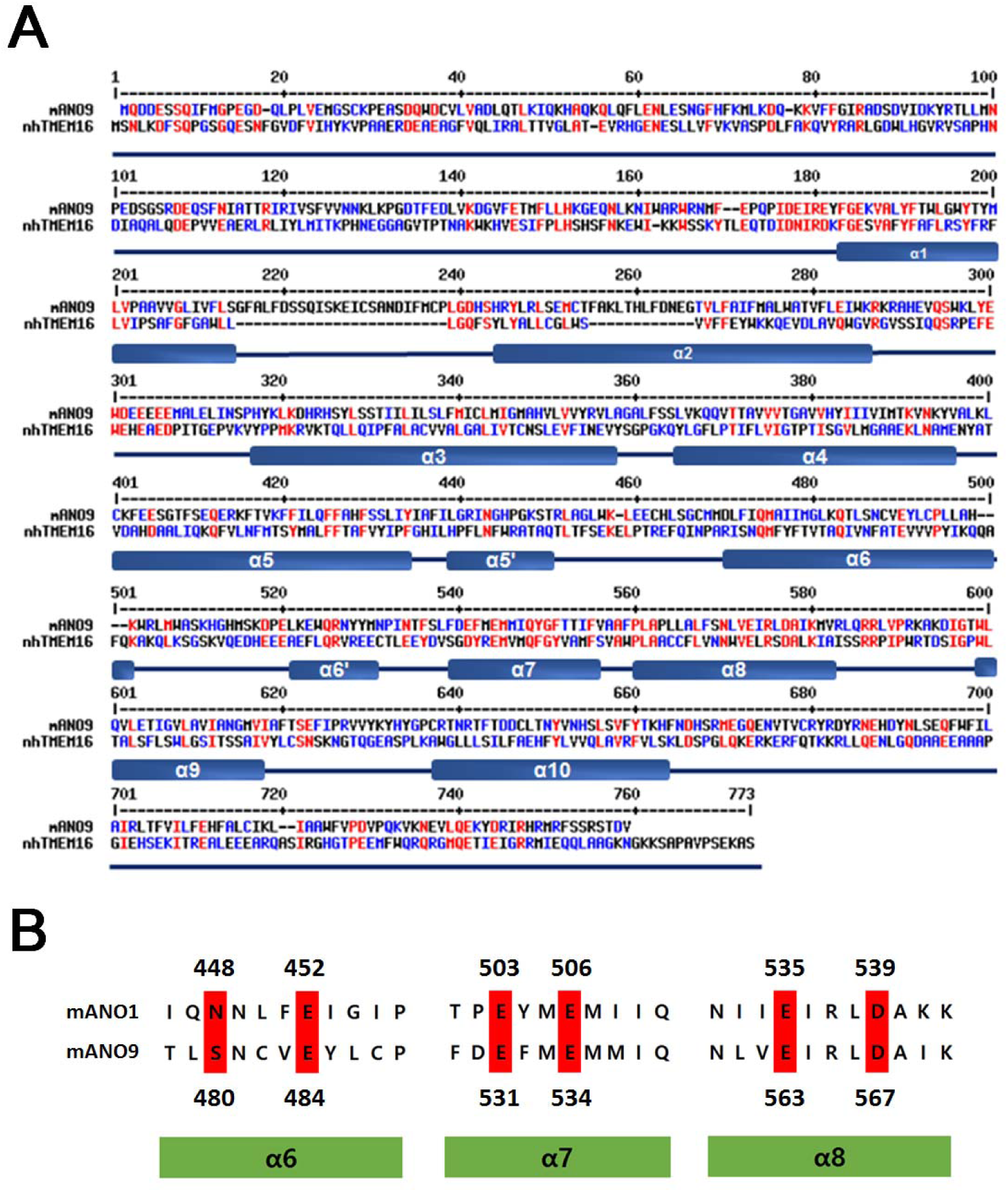
(A) Structure-based sequence alignment with mouse ANO9 and Nectria haematococca TMEM16 (nhTMEM16). High consensus residues are shown in red. α-helices in the crystal structure of nhTMEM16 were also shown in bars. (B) Sequence alignment with mouse ANO1 and mouse ANO9 at the putative Ca^2+^ binding sites in ANO1. The conserved amino acids in the putative Ca^2+^ binding sites are highlighted in red.

**Supplementary Figure 2.**
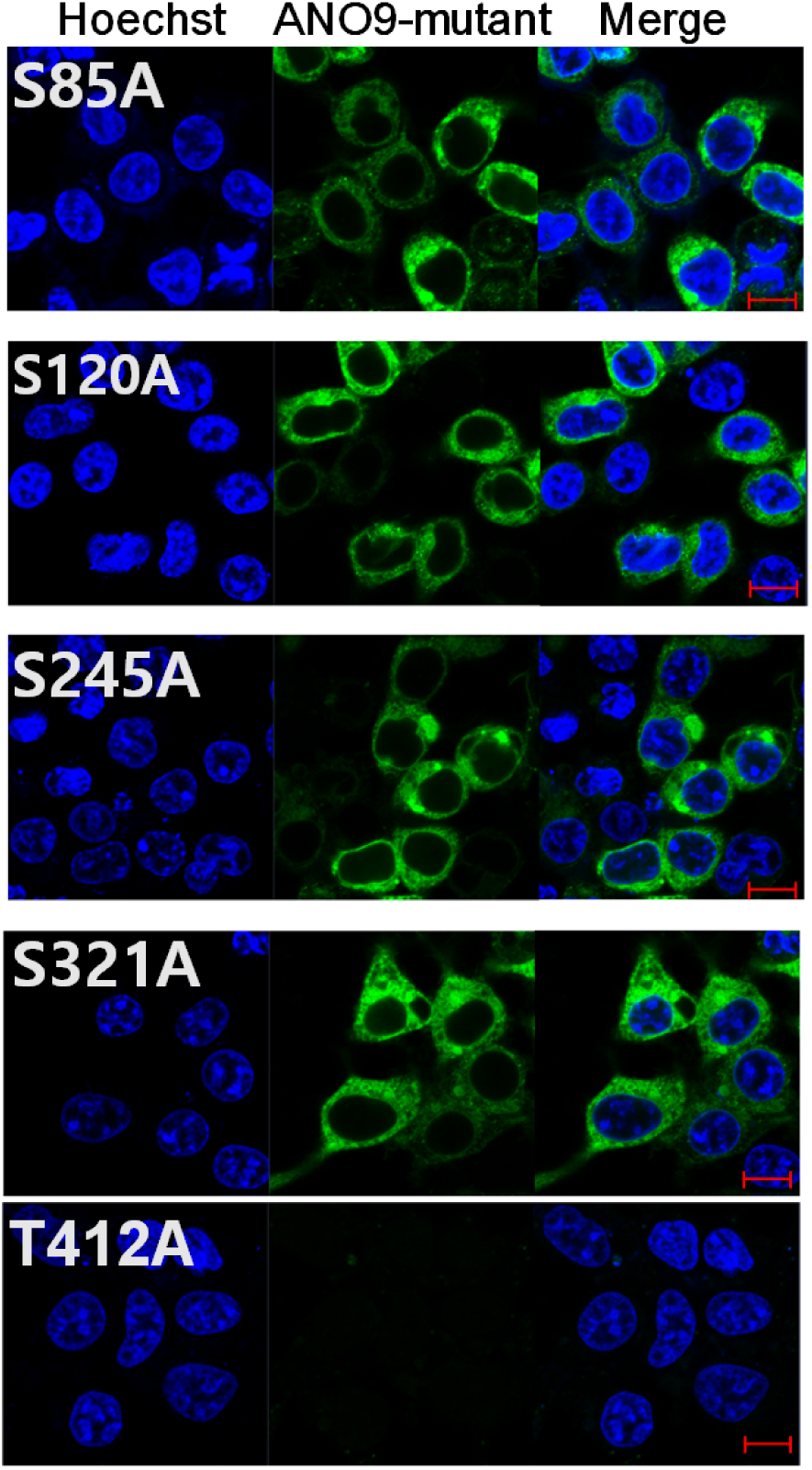
Overexpression of mANO9 mutants tagged with eGFP such as S85A, S120A, S245A, S321A, or T412A in HEK293T cells.

